# Interactions Between Hantavirus Nucleoprotein and Glycoproteins: a quantitative fluorescence microscopy study

**DOI:** 10.1101/2025.05.28.656537

**Authors:** Amit Koikkarah Aji, Salvatore Chiantia

## Abstract

Orthohantaviruses (HV) are tri-segmented negative-sense RNA viruses that can occasionally cause severe pathologies in humans. Currently, limited information exists on the molecular interactions driving HV assembly in infected cells. Specifically, it is not clear how its glycoproteins (Gn and Gc) interact with other viral or host molecules. In this study, we use one- and two-color Number & Brightness fluorescence microscopy approaches to quantitatively characterize the interactions between HV glycoproteins and the nucleoprotein NP in transfected cells. Our results indicate that HV NP homo-interactions are strongly affected by the host environment. Furthermore, we report evidence of Gc-NP interactions, based on i) the high fluorescence cross-correlation between these two proteins and ii) the increased Gc-Gc interactions observed in the presence of NP. Finally, experiments on a Gc deletion mutant suggest that the observed protein-protein interactions are mediated by the cytoplasmic tail of Gc. In conclusion, this study provides new insights into the role of the interactions between HV glycoproteins and NP in the context of HV assembly.

## INTRODUCTION

Orthohantaviruses (HVs) (Family: Hantaviridae; Order: Bunyavirales) belong to an emerging group of zoonotic viruses that cause sporadic outbreaks around the world [1]. HVs are tri-segmented negative-sense RNA viruses: the S segment encodes the nucleocapsid protein (NP), the M segment encodes the glycoproteins (GPs), and the L segment encodes the RNA-dependent RNA polymerase (RdRp) [2]. The interplay between HV GPs (named Gn and Gc) to form the outer envelope spike, consisting of Gn-Gc hetero-octamers, is the initial step of the viral assembly process [2–4]. However, limited information exists on the precise mechanisms regulating the molecular interactions involved in the subsequent stages of viral assembly. In this regard, a key event is the association between RNPs (comprised of viral RdRP, NP and viral RNA) [5] and GP spikes. This interaction is critical for viral genome packaging and influences the progression of the viral assembly by bringing pre-assembled GP spikes into proximity, eventually leading to the formation of a new virion [2].

Our current understanding of the interaction between GP spikes and RNPs in the context of virion assembly is described by the model proposed by Hepojoki et al. [2]. This states that the cytoplasmic tail (CT) of Gn initially associates with the RNP after GP spike formation. Such interaction alters the orientation of the Gn CT, thus allowing Gc to also bind to the RNP via its CT. The RNP-Gn complex further regulates viral transcription and eventually promotes viral assembly. Previous biochemical studies have attempted to validate this model by primarily identifying the interacting domains between the RNP complex and GP spikes. First, the regions within the CTs of Gn and Gc responsible for the binding to NP in multiple HV strains were mapped [6, 7]. Further, it was reported that the zinc finger domains in the Gn CT and the middle domain of NP are needed for NP-Gn interactions [7]. Additionally, the NP residues and nucleotides from genomic RNA that facilitate the binding of RNP to GP CTs were also characterized [8]. Finally, the observation of colocalization of Hantaan virus (HTNV) NP and GPs in transfected cells indicates that NP promotes Gc partition into the Golgi apparatus and that NP-Gc interactions are essential for the stabilization of the GP spike complex [9].

Despite these significant advances, some key mechanistic aspects of the proposed assembly model have not yet been explored quantitatively beyond the GP spike formation step. In this context, the molecular details of the association between NP and individual GPs are particularly relevant. HVs lack a matrix protein that mediates the interaction between the GP spike and the RNP complex and organizes the assembly of the progeny virus. The direct NP-GP association is in fact hypothesized to act as a substitute for the lack of matrix protein [10]. However, details regarding the affinity between NP and individual GPs and the role of other viral components in stabilizing the NP-GP complex remain unclear. This is primarily due to limited quantitative information on NP-GP interactions in HV assembly, particularly within living cells. Such insights are essential for accurately understanding how the viral components are packaged to form the final structure of the nascent virion.

Here, we present a quantitative characterization of NP-GP interactions using Puumala orthohantavirus (PUUV) constructs expressed in multiple living cell models. Recently, we showed significant co-localization between fluorescently labelled PUUV NP and GPs upon co-expression of NP, Gn and Gc [11]. However, these findings could only qualitatively suggest the presence of direct NP-Gc and NP-Gn interactions. In this new work, we examine NP-GP association by assessing the variations in protein multimerization and quantifying protein– protein interactions (PPI), upon co-expression of fluorescently labeled PUUV NP and GPs (Gn and Gc) in living cells. This is achieved by applying Number and Brightness (N&B) and cross-correlation N&B (ccN&B) approaches, with single cell resolution. These methods based on the statistical analysis of fluorescence fluctuations are particularly sensitive at very low protein expression levels (between µM and nM) and can assess protein behavior in a concentration dependent manner [12]. We have previously applied such techniques to analyze the role of specific PPIs in viral assembly [13] [4].

NP-GP interactions were examined in this study using different human and rodent epithelial cell models, since the pathogenesis of HV infections is influenced by the identity of the host [14]. Chinese hamster ovary (CHO) and Baby Hamster Kidney (BHK-21) rodent cells were chosen since they were previously used for the characterization of VLP formation [15], GP multimerization [4] [16], NP multimerization [17] and production of antibodies against NWHV infection [18]. Human embryonic kidney (HEK-293T) and adenocarcinomic human alveolar basal epithelial (A549) cell models were used in studies regarding host regulation factors and NP expression [19] [20], cellular immunity in infection [21] [22] and lentiviral expression of NP [23].

Our results indicate clear differences in the homotypic NP-NP interactions observed either in human or rodent epithelial cell models. Furthermore, we quantified GP multimerization and concluded that only Gc-Gc interactions are enhanced in the presence of NP. Together, these results shed light on the role of NP-GP interactions in the context of HV assembly in living cells.

## MATERIALS AND METHODS

### Cloning and generation of chimeric proteins

The constructs used for the transcription and translation of PUUV proteins originate from the PUUV Sotkamo/V-2969/81 strain. Plasmids encoding PUUV SP-FP-Gn and PUUV SP-FP-Gc, (SP: signal peptide sequence present at the N terminus of PUUV glycoprotein precursor (GPC); FP: fluorescent protein) were previously described [4, 16]. In brief, PUUV SP was introduced at the N-terminus of FP-GP (GP: Glycoprotein) to ensure physiological membrane incorporation and localization of the GP constructs [16]. The plasmid encoding PUUV YFP-NP has been previously described [11]. The construct mTurqoise2 (mTurq2)-NP was designed by substituting the mEYFP of the PUUV YFP-NP plasmid with the mTurq2. The constructs SP-mEGFP-GPΔCT and SP-mCh2-GPΔCT were produced by PCR amplification of the GPΔCT sequence (GP devoid of the cytoplasmic tail) and the subsequent product inserted between the restriction sites AgeI and Bsp1407I within the PUUV SP-FP plasmid (details on the design of PUUV FP-GP, see [4]. GnΔCT and GcΔCT sequences are described by [8] [24] respectively. All sequences were verified using the Sanger sequencing facility provided by LGC (LGC, Biosearch Technologies, Berlin).

All SP-FP-GP constructs, where the FP can be mEGFP/mEYFP/mCh2/mTurq2, is referred to as FP-GP. Any deviation from this nomenclature is explicitly highlighted in the text.

### Cell Culture and transfection

Chinese hamster ovary cells CHO-K1 (ATCC, CCL-61™), Baby hamster kidney cells BHK-21 (ATCC, CCL-10™) adenocarcinomic human alveolar basal epithelial cells A549 (ATCC, CCL-185™) and human embryonic kidney epithelial cells (HEK) from the 293T line (CRL-3216™) were purchased from ATCC, Kielpin Lomianki, Poland, and maintained in Dulbecco’s modified Eagle’s medium containing 10% fetal bovine serum, 100 U/ml penicillin, 0.1 mg/ml streptomycin, and 4 mM L-glutamine at 37°C and 5% CO_2_. Cells were passaged every 3 to 5 days, no more than 15 times. All solutions, buffers, and media used for cell culture were purchased from PAN-Biotech (Aidenbach, Germany). Cells (3 to 6 x 10^5^) were plated on glass-bottom 35-mm-diameter plates (CellVis, Mountain View, CA or MatTek Corp., Ashland, MA) 48 h before experiments. Fusion protein expression plasmids were transfected into (70 to 90% confluent) CHO and HEK 293T cells using Turbofect (Thermo Fisher Scientific, Vilnius, Lithuania) according to the manufacturer’s protocol, 20 to 24 h prior to experiments. BHK-21 and A549 cells were transfected with Lipofectamine 3000, according to the manufacturer’s protocol (Thermo Fisher Scientific, Vilnius, Lithuania), 20 to 24 h prior to experiments.

### Confocal microscopy imaging

3 to 6 x 10^5^ cells were plated onto 35-mm glass-bottom dishes (CellVis, Mountain View, CA or MatTek Corp., Ashland, MA) 48 h prior to the experiment and transfected 16 to 24 h post seeding. Confocal images were acquired on a Zeiss LSM780 system (Carl Zeiss, Oberkochen, Germany) using a 40x DIC M27 water immersion objective. To limit the out-of-focus light, a pinhole with size corresponding to one Airy unit (∼39 μm) was used. Samples were excited with a 488-nm argon laser and a 561-nm diode laser. The fluorescence signal was collected by a Zeiss QUASAR multichannel GaAsP detector in photon-counting mode. The laser power was set so that the photon count rate and bleaching remained below 1 MHz and ca. 20%, respectively (typically ∼3 μW for 488 nm, and ∼5 µW for 561 nm). Measurements exhibiting stronger bleaching were discarded. Fluorescence was detected between 499 and 552 nm (mEGFP, Alexa Fluor 488) and between 570 and 695 nm (mCh2, Alexa Fluor 568), after passing through an MBS 488/561 dichroic mirror.

### Number and brightness (N&B)

Number and brightness analysis was performed as previously described by [4]. Briefly, images of 128 x128 pixels were acquired with pixel dimensions of 400 nm and a pixel dwell time between 25 and 50 µs. Image time-stacks of 100 scans were collected using the Zeiss Blu ZEN software. The intensity time-stacks data were imported into MATLAB using the Bioformat [25] package and analyzed using a self-written code (The MathWorks, Natick, MA, USA). The algorithm uses the equations from [26] for the specific case of photon-counting detectors to obtain the molecular brightness and number as a function of pixel position. Upon selecting the required region of interest (ROI), fluorophore bleaching and minor cell movements within a few pixels are suitably corrected. ROIs did purposedly not include any immobile structures, such as large aggregates or immobile protein filaments. Pixel intensities are then used to calculate the average brightness of the FP detected within the cellular ROI. This average brightness is utilized to evaluate the multimerization state of the FP upon comparison to a reference monomer brightness [27]. Thus, the oligomeric size of the protein complex is evaluated in terms of monomer units.

### Cross Correlation Number and Brightness (ccN&B)

The ccN&B analysis is based on the workflow described by [13] and modified suitably for the evaluation within the cytosolic region of the cell. Briefly, an image stack is acquired over time typically consisting of 100 frames. Images of 256 x y (<u>< 256)</u> pixels with a pixel size of 0.21 µm and a pixel dwell time of 12.6 µs were acquired alternating two different excitation wavelengths. CZI image output files were imported into MATLAB using the Bioformat [25] package and analyzed using a self-written script implementing the framework described by [28] for the specific case of photon-counting detectors and two-color excitation. Pixels corresponding to a ROI are manually selected in an image map. Next, to account for lateral drift during the acquisition, frames are aligned to the first frame by maximizing the spatial correlation between sub-selections in consecutive frames, averaged over both channels, as a function of arbitrary translations. Additional steps regarding corrections for bleaching, minor cell movements & specific detector response are applied as described in [13].

The ccN&B analysis calculates the dimensionless parameter relative cross correlation (Relative CC). Relative CC is calculated as described by [13]:

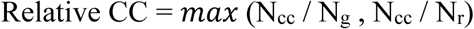

Here, N_g_ and N_r_ are the particle numbers detected in the image channel corresponding to the green FP and red FP, respectively. N_cc_ is obtained by dividing the product of the average intensities of the green and red FP channels by the covariance of the two channels.

### Statistical Analysis

The statistical analysis in this work was performed using GraphPad Prism version 9.0.0 (GraphPad Software, LCC, San Diego, CA, USA). For pairwise comparisons of the derived parameters, Kruskal Wallis Test and modified Tukey’s multiple comparisons test were used.

## RESULTS

### NP-NP interactions in human epithelial cells are stronger than in rodent epithelial cells

Hägele et al. [29] observed significant variations in NP localization and function between infected Vero E6 cells and primary human renal cells. This allowed them to conclude that the behavior of NP might depend on the specific cell type. Here, we aimed to determine whether different host environments (namely, human vs. rodent epithelial cell models) might influence homotypic NP-NP interactions.

NP multimerization in cells imaged via confocal microscopy was examined using N&B analysis, a quantitative approach based on the statistical fluctuations of the fluorescence signal within the probed confocal volume. This approach was previously used to monitor NP multimerization in CHO cells [11]. In this work, we extend the analysis to compare the average NP multimerization detected in human epithelial cells (HEK and A549) and rodent epithelial cell models (BHK-21) 16 to 20 hpt.

As previously shown Davies et al. [30], NP initially appears homogenously distributed in the cytoplasm. This is followed by the formation of small mobile clusters, for all cell types tested (Figure 1 A-D), after ca. 4 hours post transfection. At later time points (i.e. after ca. 12 to 16 hours), NP starts to coalesce forming large immobile filaments [11], but such samples were not included in the current analysis.

**Figure 1:**
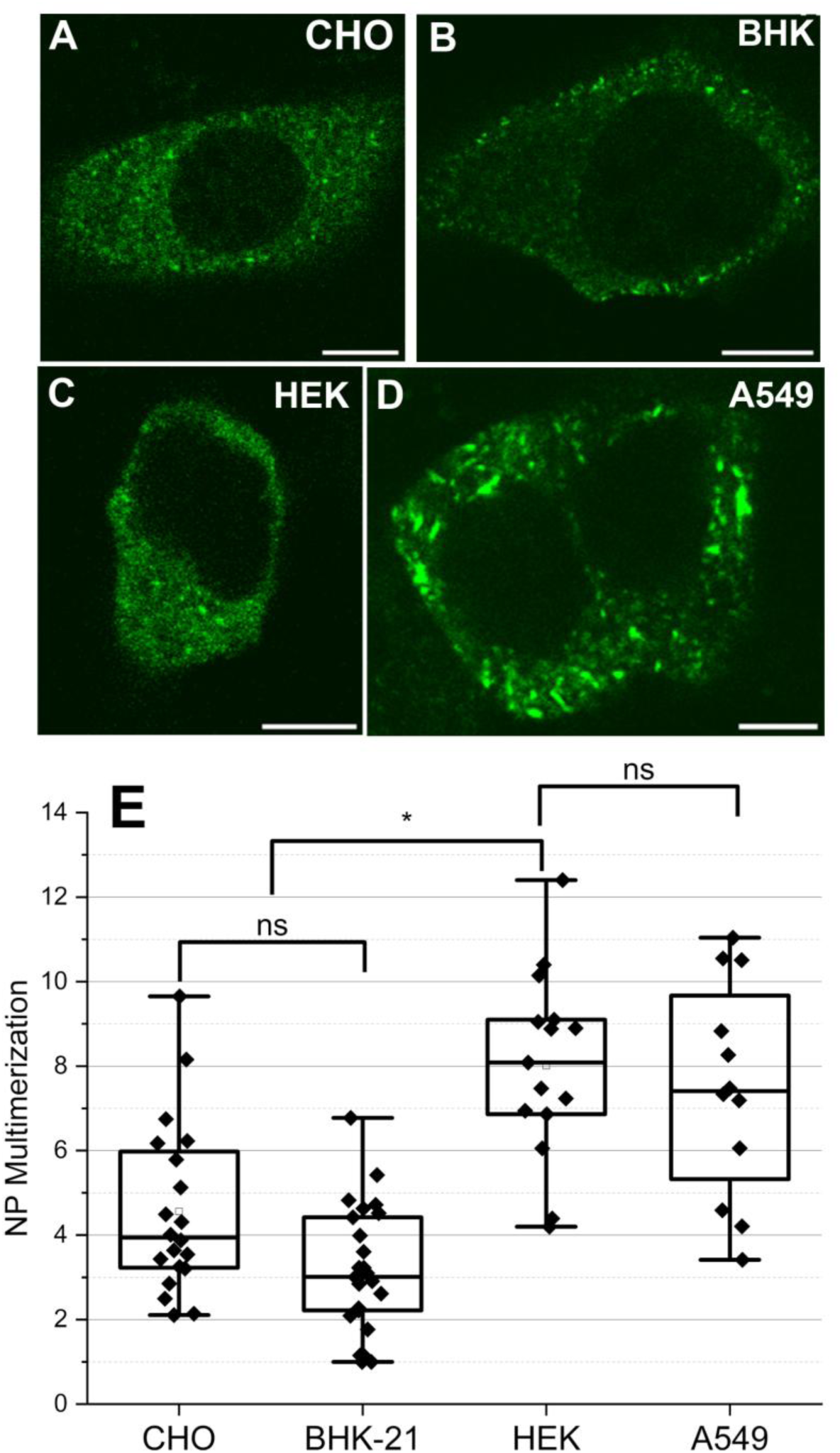
Average multimerization of fluorescently labelled PUUV NP is higher in human cell models compared to rodent epithelial cell models. YFP-NP was expressed in multiple epithelial cell models and observed 16 to 20 hpt. Panels A-D show representative confocal microscopy images of [A] CHO, [B] BHK-21, [C] HEK 293T and [D] A549 cells expressing YFP-NP. Panel E shows the quantitative comparison of YFP-NP multimerization between different cell models. For this analysis, cells expressing YFP-NP within the concentration range 0.1-0.5 µM were chosen. Data points for NP multimerization in CHO cells are from [11] and incorporated as reference values. Each point in the boxplot represents the average multimerization in one cell. At least 15 cells were analyzed for each case. Statistical analysis was performed using the pairwise Kruskal Wallis one way ANOVA test (ns: not significant, * p< 0.01). Scale bars are 10 µm.

According to N&B analysis, NP forms small oligomers, up to ca. decamers in average, for all cell models tested, within the entire concentration regime explored (Figure S1). More specifically, NP multimerization values in BHK cells increase strongly with protein concentration, as previously observed for CHO cells [11] (shown in Figure S1, as reference). On the other hand, multimerization values measured in A549 and HEK cells are higher (especially at lower concentrations) and display a less marked correlation with protein concentration (Figure S1). Since different cell models displayed variability in the expression levels, we compared the average NP multimerization in human and rodent cell models within a restricted concentration range (0.1 to 0.5 µM). Within such a concentration range, we were able to collect a similar amount of data points for all cell models. In these conditions, NP forms on average tetramers in rodent epithelial cells and roughly octamers in human epithelial cells (Figure 1E).

We further attempted a statistical analysis of the multimerization-concentration curves, for different cell models and across the whole available protein concentration range, by fitting an empirical model to the data [31], as shown in Figure S1. In agreement with the above-mentioned observations, NP multimerization behavior in rodent (CHO [11] and BHK-21) and human epithelial cell models (HEK-293T and A549) appears to be significantly different over the observed concentration range (Supplementary Table 1).

Altogether, these results indicate that the average NP-NP interactions observed in human epithelial cells are stronger than in rodent epithelial cell models, at least at the early stages of NP expression and assembly.

### NP interacts with Gc within few small intracellular regions

We proceeded to compare the interactions between PUUV NP and each PUUV GP (Gn or Gc) upon co-expression in different human (HEK 293T and A549) and rodent epithelial (CHO and BHK-21) cell models. Welke et al. [11] previously indicated that NP, Gn and Gc displayed significant spatial colocalization in transfected CHO cells. While these findings indicate that NP and GPs are found in close vicinity to each other, they do not directly imply an interaction between the proteins. Therefore, we performed two color ccN&B analysis to investigate whether NP significantly interacts with any of the GPs. This method provides a parameter (i.e., relative CC) that indicates the relative amounts of co-diffusing hetero complexes.

Figure 2A to 2C and 2F to 2H show representative confocal microscopy images of cells co-expressing YFP-NP and either mCh2-Gc or mCh2-Gn in CHO cells, respectively. The intracellular localization of Gn and Gc within the ER and/or perinuclear region is similar to what was previously observed [4]. To quantify hetero-interactions, ccN&B analysis was performed, thus obtaining relative CC maps, as shown in Figure 2 D and 2 E for NP-Gc and NP-Gn interactions, respectively. Additional examples of relative CC maps are shown in Figure S3 I and J. Overall protein-protein interactions were first quantified by averaging the relative CC from each pixel in the selected ROIs (roughly corresponding to the whole intra-cellular region). As shown in Figure 2 I for CHO cells, significant interactions were detected for the positive control (i.e., a tandem cytosolic construct of YFP and mCh2) but not for the negative control (consisting of independently co-expressed YFP-NP and cytosolic mCh2). Similar results were obtained for the controls in all cell models tested (data not shown). The interactions between NP and either GP (i.e., Gn or Gc), averaged over all the whole cell, were relatively weak for all cell models tested (CHO cells, see Figure 2I; HEK cells, see Figure S2A and A549 cells see Figure S2B) and not distinguishable from the negative control. Co-expression of fluorescently labelled NP and GP constructs in BHK-21 cells did not yield a GP expression high enough for quantifying relative CC and, therefore, this cell model was not used for subsequent analysis.

**Figure 2:**
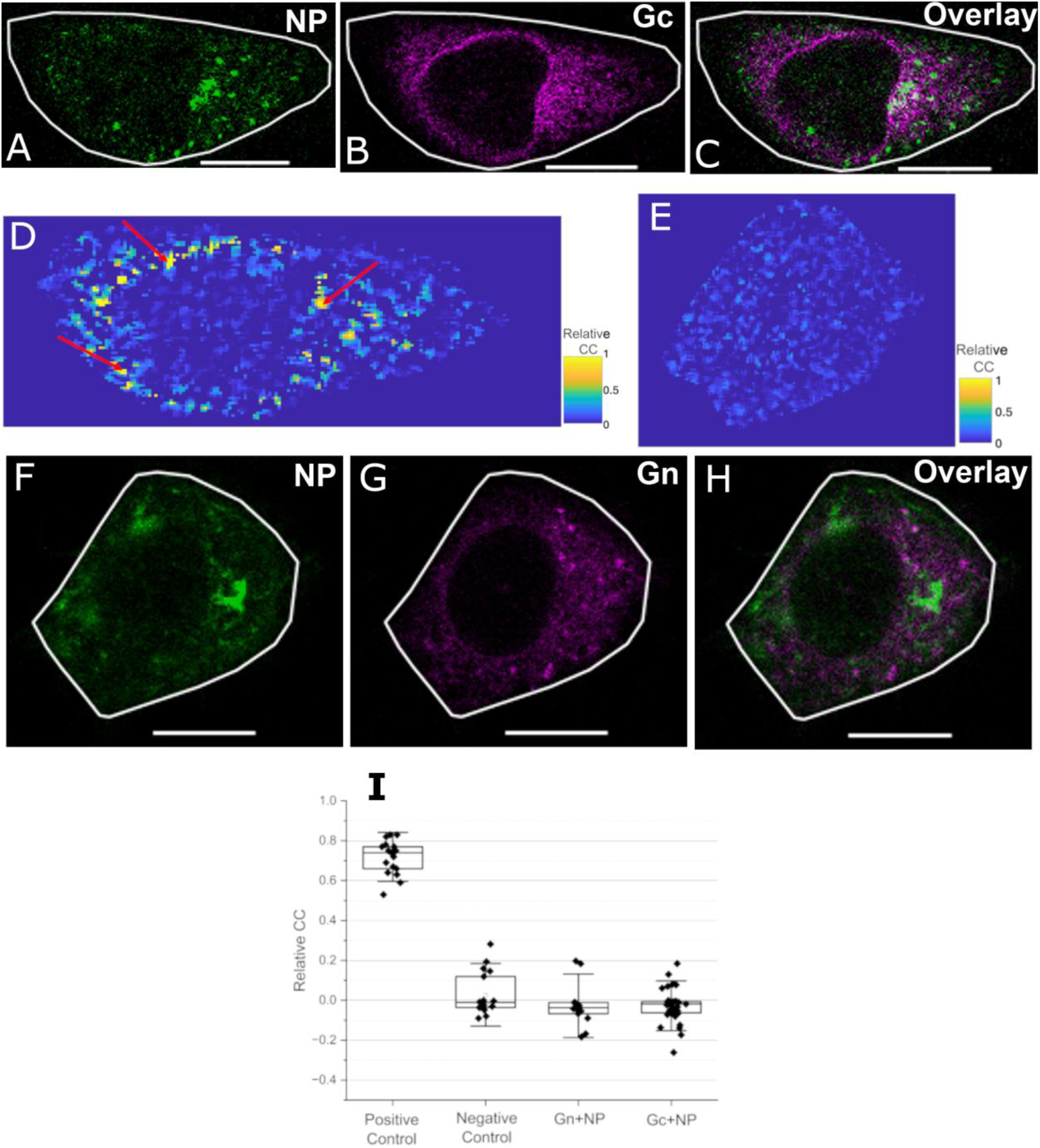
Single-cell analysis and spatial maps of the interactions between NP and GPs. **Panel**s A-C show representative images of CHO cells co-expressing PUUV YFP-NP and PUUV mCh2-Gc, 20 to 24 hpt. Panel D shows the pixel-by-pixel relative CC map of the same CHO cell, within a ROI as indicated by a white line in panels A-C. Red arrows in panel D denote localized regions characterized by high relative CC values. Panel E shows the pixel-by-pixel relative CC map of the same CHO cell, within a ROI as indicated by a white line in panels F-H. Panels F-H show representative images of CHO cells co-expressing PUUV YFP-NP and PUUV mCh2-Gn, 20 to 24 hpt. Panel I shows a box plot of the relative CC values averaged over whole-cell-ROIs and measured for different constructs expressed in CHO cells. Positive control is a tandem YFP-mCh2 construct that localizes in the cytoplasm. The negative control refers to cells co-expressing YFP-NP and cytosolic mCherry2. ROIs in cells with intensity values less than 1MHz in the YFP and mCherry2 channel were chosen for evaluating the average relative CC within each cell. Each point in panel I represents the average relative CC value from one ROI in one cell. Number of points for each case > 15, from 3 independent experiments. Scale bars are 5 µm.

Next, we analyzed more in detail the spatial variations of the relative CC parameter in cells expressing the different constructs. As expected, CHO cells expressing the tandem mEYFP-mCh2 construct (positive control) exhibited positive relative CC values homogenously distributed throughout the analyzed intracellular ROI (Figure S3 E-H). Similarly, the analysis of CHO cells expressing YFP-NP and cytosolic mCherry2 (negative control) resulted in homogeneous low relative CC values throughout the cell (Figure S3 A-D). Interestingly, similarly to the negative control, we observed relatively homogeneous intracellular distribution of low relative CC values for cells co-expressing Gn and NP (Figure 2 E and S3 I). On the other hand, we consistently noticed localized regions characterized by high relative CC values (i.e., high local abundance of co-diffusing Gc-NP complexes) in CHO cells co-expressing Gc and NP (see red arrows in Figures 2 D and S3 J). Such regions were mostly observed around the perinuclear region and, most importantly, do not display an evident correlation with the spatial distribution of either Gc or NP (see Figure 2 A, B and D). Similar results were obtained in other cell models (data not shown).

Altogether, these results demonstrate that PUUV GPs interact in average weakly with PUUV NP in intracellular regions, for all cell models tested. Nevertheless, PUUV Gc appears to interact significantly with PUUV NP within localized intracellular regions close to the perinuclear region.

### Apparent Gc multimerization increases in the presence of NP

The experiments described above suggest that NP might play a role in intermolecular interactions involving Gc, but not Gn. To further explore such interaction network, we measured GP-GP interactions (i.e. the multimerization state of each GP) in the presence of NP, using single color N&B in different cell models. As for the ccN&B analysis, cells displaying a quasi-homogeneous distribution of NP within the cytoplasm (see Figure 1 A-D) were chosen.

PUUV Gc multimerization in the presence of PUUV NP was analyzed by co-expressing YFP-NP and mCh2-Gc in CHO (Figure 2 A - C), HEK (Figure S4 A-C) and A549 (Figure S4 D-F) cells and performing N&B measurements 24 hpt. Our previous work shows that expression of Gc alone results in a mixture of monomers and dimers in the ER of different cell models [4] [16]. Strikingly, in the presence of NP, Gc forms instead large multimers (ca. up to octamers) for all the cell models tested, within the explored concentration range (Figure 3A). Thus, Gc-Gc interactions in the presence of NP are significantly stronger than those previously observed in the absence of NP (shown in black in Figure 3 A as reference, [4].

**Figure 3:**
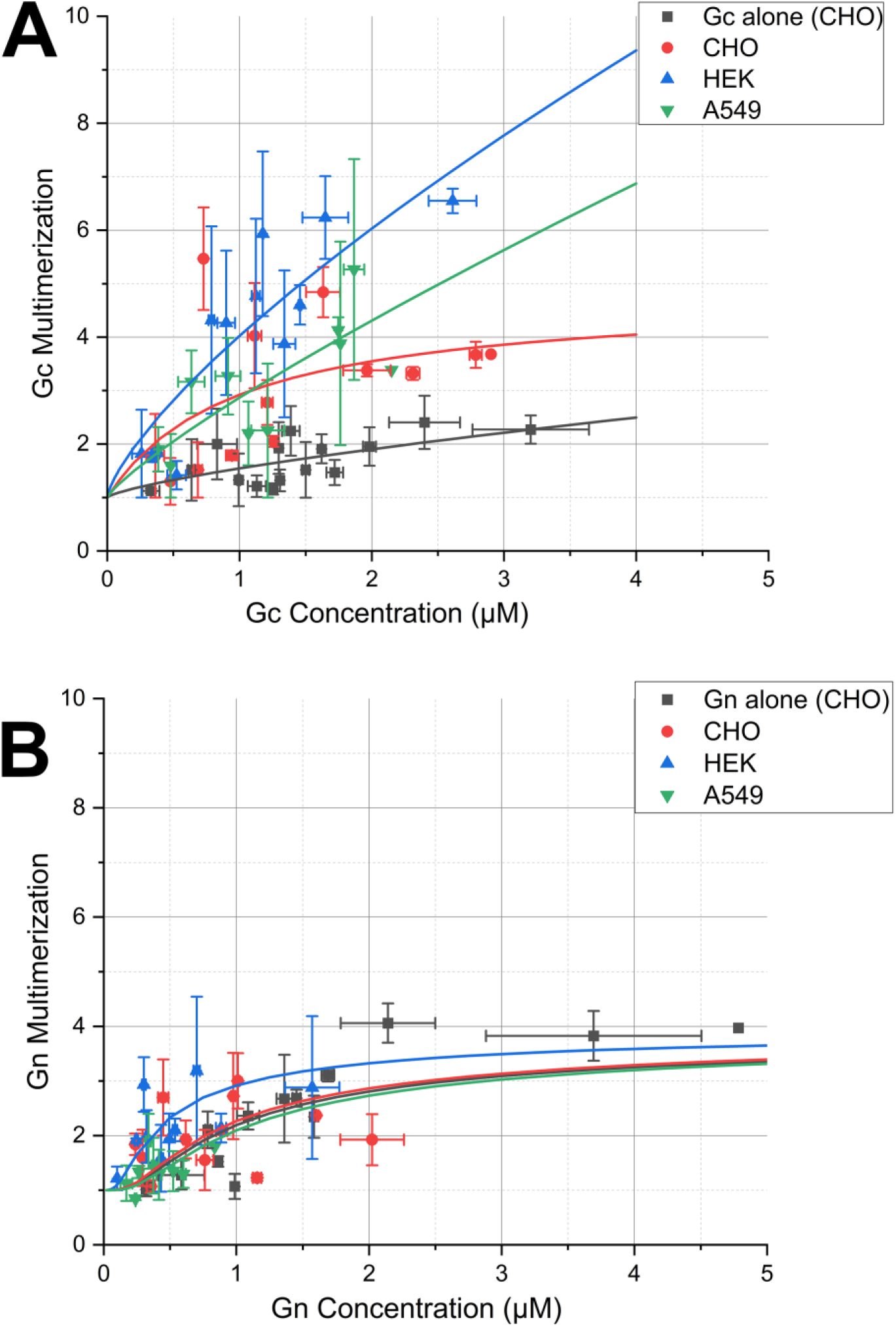
PUUV Gc (but not Gn) multimerization behavior is altered in the presence of NP. Panel A shows the concentration-dependent multimerization analysis of PUUV Gc in the presence of PUUV NP using N&B, in different epithelial cell models: CHO (red), A549 (green) and HEK 293T (dark green). PUUV Gc multimerization in the presence of PUUV NP is compared to that measured in the absence of NP in CHO cells (black, data from [4]. Each point in the graph represents the binned average multimerization values from two cells. The solid lines represent a fit to an empirical power growth model [31] in the form of y= 1+ a*x^k^. Fit results and statistical analysis are shown in Table S2. Panel B shows the concentration-dependent multimerization analysis of PUUV Gn in the presence of PUUV NP using N&B, in the same epithelial cell models as for panel A. PUUV Gn multimerization in the presence of PUUV NP is compared to that measured in the absence of NP in CHO cells (black, data from [4]. Each point in the graph represents the binned average multimerization values from two cells. The solid lines represent a fit to a monomer-tetramer equilibrium model [32]. Fit results and statistical analysis are shown in Table S3.

To compare different datasets, the multimerization curves were analyzed using a simple empirical growth model with two free parameters [32]. The resulting curves for all the examined cells expressing both Gc and NP lay considerably above the multimerization curve obtained in CHO cells expressing only Gc (p<0.01), thereby confirming the initial qualitative observation. Only minor differences can be observed between the other curves, mostly due to the large data spread (see Table S2).

### Gn remains in a monomer-tetramer equilibrium, independent of the presence of NP

We further examined the multimerization of Gn in the presence of NP. CHO (Figure 2 E-G), HEK (Figure S5 A-C) and A549 (Figure S5 D-F) cells co-expressing fluorescently labelled Gn and NP were observed 20 to 24 hpt. Cells/ROIs in which NP formed homogeneously distributed small mobile oligomers were selected, thus excluding from the analysis large immobile NP structures. We have previously shown that Gn forms up to tetramers in the absence of other viral proteins [4]. The multimerization values obtained in this work (Figure 3 B) in the presence of NP are compatible with a similar multimerization behavior, for all the cell models tested, at least within the restricted concentration range that could be explored.

To quantitatively compare the different datasets, multimerization curves were fitted to an analytical model that describes a monomer to tetramer equilibrium through the association constant K_4_ [32]. This parameter was not statistically distinguishable among the different samples (0.2<p<0.8) (see Table S3). This result indicates that Gn-Gn interactions are not strongly affected by the presence of NP.

### Deletion of Gc-CT domain abolishes the enhancement of Gc-Gc interactions induced by NP

Finally, the influence of the GP-CT domain on GP multimerization in the presence of NP was examined, since NP was proposed to interact with GPs via their CTs. Truncated GPs without CT domains (GPΔCT) were initially expressed in cells in the presence and absence of NP (Figure 4 A-D for GcΔCT, Figure S6 A-D for GnΔCT). GPΔCT multimerization was then analyzed 20 to 24 hpt. Figure 4E shows that fluorescently labelled GcΔCT, in the absence of NP, displays an average multimerization of ∼1.4 at the highest measured concentration. According to this value and assuming a monomer-dimer equilibrium, the protein appears to form mostly monomers (ca. 75 mol%), in agreement with previous measurements on non-fluorescent constructs *in vitro* [33]. In the presence of YFP-NP, the multimerization of GcΔCT did not change substantially (Figure 4 E). Similar results were obtained with A549 cells (Figure S7). The concentration dependent multimerization data points in Figure 4 E were analyzed with an analytic model describing monomer-dimer equilibrium [32]. The derived association constant K_2_ did not change significantly in the presence or absence of NP (Supplementary Table 4).

**Figure 4:**
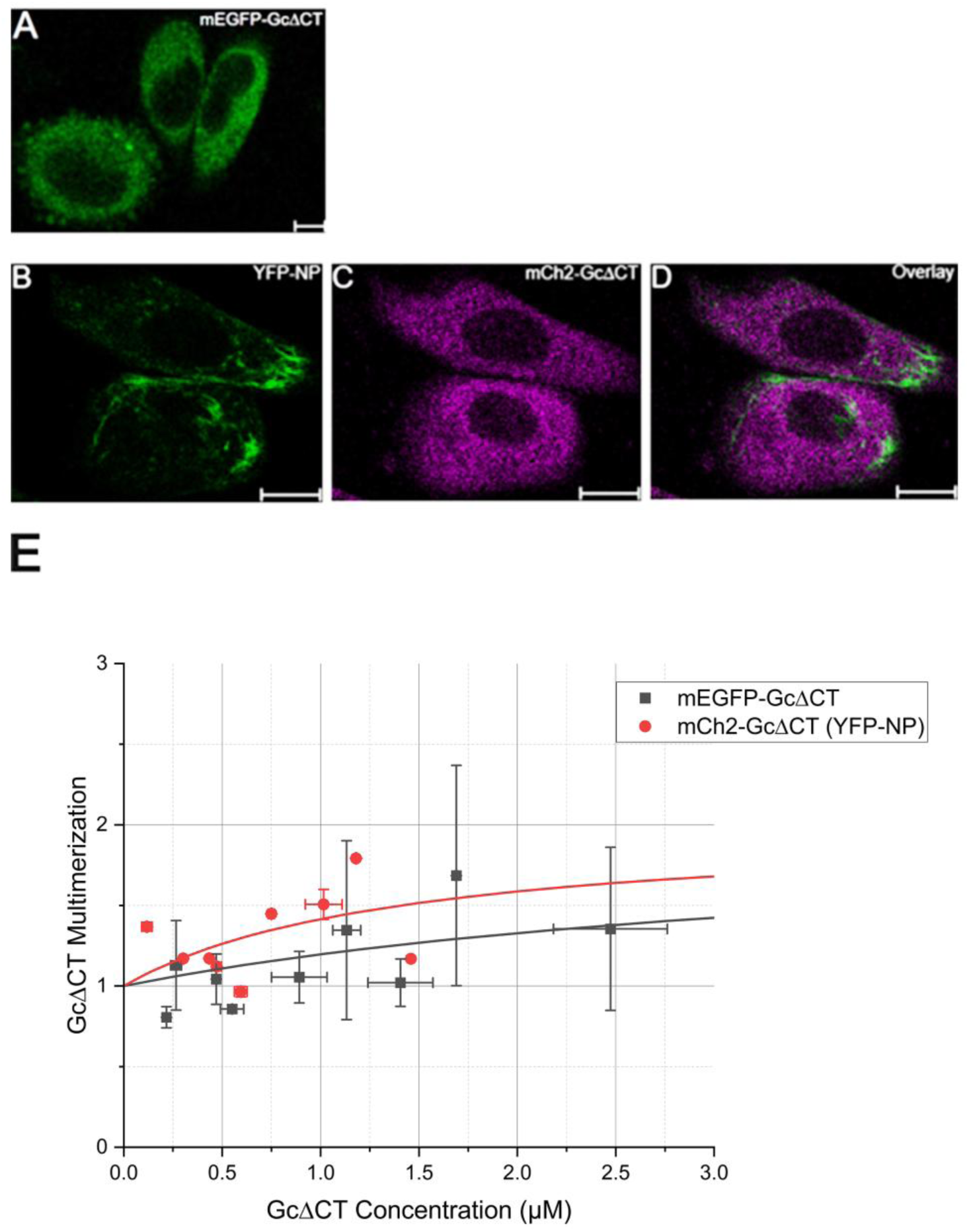
Deletion of GcΔCT inhibits the formation of large Gc assemblies in the presence of NP. Panel A shows a representative image of CHO cells expressing PUUV mEGFP-GcΔCT and observed 24 hpt. Panels B - D show typical confocal image of CHO cells co-expressing YFP-NP and mCh2-GcΔCT. Panel E shows the concentration-dependent multimerization analysis of PUUV GcΔCT in the absence (black) and presence (red) of PUUV NP, calculated using N&B analysis. Each point in the graph represents the binned average multimerization from two cells. The solid lines represent a fit to a monomer-dimer equilibrium model [32]. Fit results and statistical analysis are shown in Table S4. Scale bars are 10 µm.

The influence of Gn-CT on Gn-Gn interactions was similarly analyzed. Figure S6 E shows that fluorescently labelled GnΔCT, in the absence of NP, displays an average multimerization value compatible with the presence of a large number of monomers in equilibrium with dimers, in agreement with the results obtained using non-fluorescent constructs [15]. In the presence of YFP-NP, the average multimerization of GnΔCT remained similar. Analogous results were obtained with A549 cells (Figure S8).

In conclusion, the increased formation of Gc-Gc complexes induced by NP appears to be mediated by the Gc-CT domain, for all the cell models tested. On the other hand, we did not observe any influence of NP on the multimerization of Gn, irrespective of the presence of its CT.

## DISCUSSION

NP is a multifunctional HV protein that is capable to interact with different host proteins, other viral components and mediates the association between the RNP and GPs ([34]. While previous studies focused on the protein regions possibly involved in the binding between NP and GPs, there is limited information regarding the molecular details of the actual interactions between NP and either Gc or Gn occurring *in vivo*. In this study, we address this important question by quantifying the interaction between NP and GPs upon co-expression, at the single cell level. In order to pinpoint specific interactions between NP and Gn or Gc, we evaluated the behavior of these proteins in the absence of other viral components (e.g. viral RNA) in live cell models, through transient transfection.

HV infections demonstrate distinct cellular tropism [14] [21] [29]. Therefore, we initially assessed if a specific cellular environment could influence NP-NP binding, by comparing human and rodent epithelial cell models. We observed that average homotypic NP-NP interactions are significantly stronger in human epithelial cell models compared to rodent epithelial cell models, at a given time point post-transfection. This observation is in line with the results of [29], who reported point-like NP structures in infected Vero E6 cells, smaller than the filamentous NP assemblies present instead in infected human renal cells. Also, it has been shown that cytoskeletal components, such as actin, vimentin and microtubules, influence NP organization in a fashion that varies among different HV strains and cell models [29] [35]. It is therefore possible that the modulation of NP-NP interactions described in this study may be determined by cell type-specific interactions with cytoskeletal components. Alternatively, post-translational modifications, including phosphorylation [36] [37] or SUMOylation [38] may also contribute to the regulation of NP multimerization in a cell type-dependent manner. A third possibility is that NP multimerization is influenced by other host-specific factors [39], e.g. proteins binding to the NP multimerization interface. Independent from the specific molecular mechanism, it would be very interesting to investigate whether alterations in NP multimerization equilibrium are truly correlated to the susceptibility of a specific host cell to HV infection.

In the second part of our investigation, we observed weak overall NP-GP interactions, independent of the tested cell model, while NP still mostly formed small mobile clusters. Of interest, within single cells, localized regions were detected in which NP showed an apparent strong interaction with Gc. According to our data, such regions do not correspond to locally increased protein concentration (data not shown). We are currently investigating whether locally enhanced NP-Gc interactions might instead exhibit a spatial correlation with e.g. cytoskeletal elements or other intracellular structures. Interestingly, such local interactions with NP were never observed for Gn. Previous studies have shown significant interactions between these two proteins via Gn CT domain in infected cells using co-immunoprecipitation [6]. Nevertheless, it must be noted that these experiments were performed several days post infections (i.e., at a time point when NP forms large immobile structures [30] and in the presence of other viral components. This observation suggests that NP-Gn interactions might be possibly mediated by viral RNA [6] [8]. Also, our results additionally show that the quantitative microscopy approach with single-cell resolution described here can provide more detailed information, compared to bulk methods that average over a large number of cells.

While the fluorescence fluctuation microscopy analysis used in this work (NaB) delivers quantitative information about inter-molecular interactions and has a high spatial resolution, it also requires the observed molecules to be mobile during the observation. Although we restricted our analysis to cells which did not show large immobile structures, it is known that NP might form a small fraction of larger very slow NP aggregates, as described by [11]. If the reported higher CC values (i.e. stronger interactions) are indeed originating from such almost immobile NP fractions, NP-GP interactions cannot be quantified precisely with this approach. For this reason, we complemented these measurements by separately observing GP multimerization in the presence on NP.

In the case of Gc, we observed a striking difference in homotypic interactions when NP is also present. While not a direct measurement of NP-Gc interaction, this is compatible with the above-mentioned results, under the hypothesis that the observed Gc multimer formation is indeed somehow mediated by interactions with NP (e.g., through the binding of several individual Gc molecules to a large NP multimer). Gc localization within NP clusters was also previously observed for other Bunyaviruses, such as Tomato spotted wilt virus (TSWV) [40] [41].

Furthermore, our experiments on truncated Gc constructs indicate that NP-Gc interaction occurs via Gc CT domain, as also suggested by previous studies [2] [9]. Alternatively, it is also possible that the removal of the CT affects the stability and structure of Gc [16] and, thus, its interaction with NP. Taken together, our findings suggest that HV NP can form complexes with Gc (possibly, including other molecules as well), also in the absence of Gn or RNPs.

On the other hand, the lack of direct interactions between Gn and NP as observed via ccNaB, is corroborated by the fact that, in the observed cell models, Gn multimerization is independent of the presence of NP, with this GP forming consistently monomers to tetramers, as expected.

In conclusion, we have investigated here the interactions between HV NP and GPs using quantitative fluorescence microscopy approaches. According to NaB experiments, NP can interact with Gc in the absence of other viral components, but not with Gn. The observed interactions appear to be mediated by the CT of Gc. Of note, all the experiments performed in this work involve the transient expression of up to two viral proteins at the same time, with the purpose of specifically identifying direct interactions (i.e. not involving other viral proteins). As mentioned above, a more comprehensive picture of the interaction network can be obtained if additional viral components are systematically added to the studied system (e.g. RNA to further characterize NP-Gn interactions). Future studies will for example focus on quantifying interactions between fluorescently labelled NP, Gn and Gc, also using third-order correlation functions [42]. Such experiments are complicated by the interference of fluorescent labels on inter-molecular interactions (e.g. between Gn and Gc) [4]. By using more advanced methodologies for labelling (see e.g. ALFA tag [43], bioorthogonal click chemistry [44], the approach described in this work open the avenues for the precise characterization of binding between other HV proteins and further elucidating the steps involved in the HV virion formation.

### SUPPLEMENTARY MATERIALS

Supplementary information is provided in the form of 4 supplementary tables and 8 supplementary figures in one single document. Numerical data and raw data (i.e., before binning) for all figures are provided in the form of two spreadsheets.

### AUTHOR CONTRIBUTIONS

Conceptualization, A.K.A. and S.C.; Methodology, A.K.A. and S.C.; Software, A.K.A. and S.C.; Validation, A.K.A. and S.C..; Formal Analysis, A.K.A.; Investigation, A.K.A.; Resources, S.C.; Data Curation, A.K.A.; Writing – Original Draft Preparation, A.K.A. and S.C.; Writing – Review & Editing, A.K.A. and S.C.; Visualization, A.K.A.; Supervision, S.C.; Project Administration, S.C.; Funding Acquisition, S.C.

## FUNDING

This work was supported by the German Research Foundation (Deutsche Forschungsgemeinschaft), grant 407961559 to S.C.

## DATA AVAILABILITY

The original contributions presented in this study are included in the supplementary material. Further inquiries can be directed to the corresponding author.

## Supporting information

Data from figures

Raw data from figures

## ACKNOWLEDGMENTS

We thank Dr. Titas Mandal for helping with the manuscript revision.

## CONFLICTS OF INTEREST

The authors declare no conflict of interest.

## Notes

### Competing Interest Statement

The authors have declared no competing interest.

