## Supplementary Materials for "Interactions Between Hantavirus Nucleoprotein and Glycoproteins: a quantitative fluorescence microscopy study"

### Supplementary Information

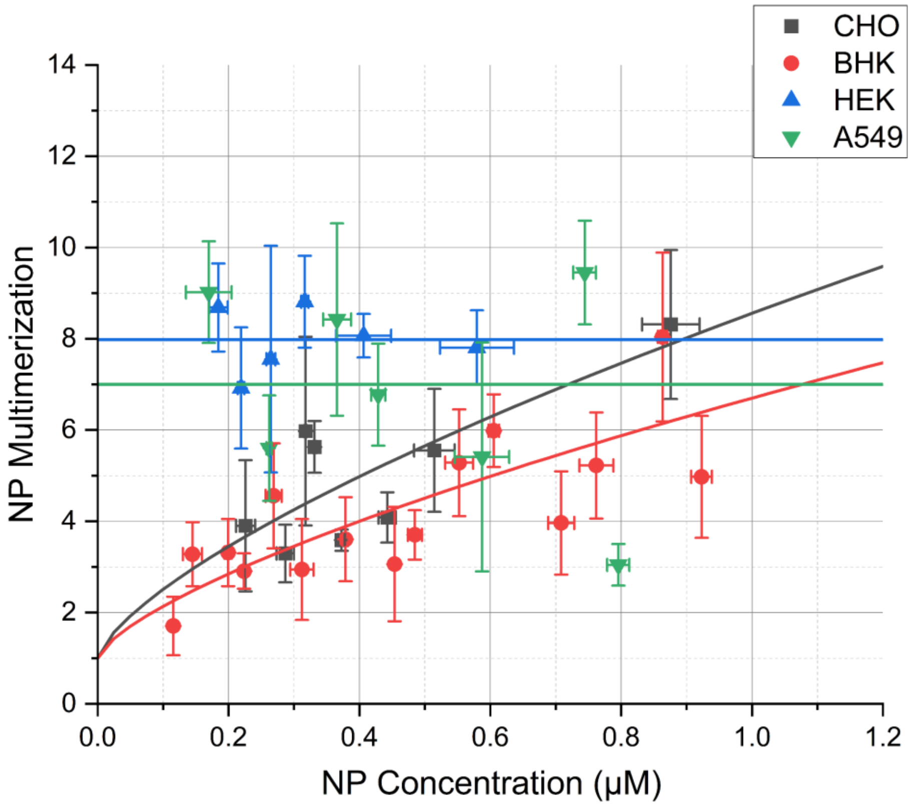


**S1: Homotypic NP-NP interactions differ between human and rodent epithelial cell models.** Concentration-dependent NP multimerization plotted for multiple epithelial cell models. An empirical multimerization model is fitted to the data to assess the differences between the cell types ([31]) ($y=1+a\cdot x^k$, continuous lines). Since NP multimerization in A549 and HEK cells is basically constant in the explored concentration range, it could only be fitted with a constant model (i.e., k=0). NP multimerization points from CHO (from [11], BHK-21, HEK-293T and A549 cell models are represented in black, red, blue and green, respectively. Each point represents the average values from 2 to 4 cells pooled together, with SD as error bars. Table S1 shows the results of the statistical analysis for the comparison of the parameters *a* and *k*, across all cell models. For all obtained curves, the parameter *a* is not statistically distinguishable ($p>0.47$, one-way Anova Tukey correction). Therefore, no significant difference can be observed between CHO and BHK cells, or HEK and A549 cells. On the other hand, the NP multimerization behavior observed in rodent epithelial cell models is significantly different from that observed in human epithelial cell models.

| Cell Line | *a* | SD | *k* | SD | N pooled | N total |
|---|---|---|---|---|---|---|
| CHO | 6.7 | 2.5 | 0.6 | 0.3 | 8 | 23 |
| BHK | 5.2 | 0.7 | 0.60 | 0.10 | 15 | 45 |
| HEK | 7 | 3.0 | 0, fixed | - | 9 | 18 |
| A549 | 6.6 | 1.5 | 0, fixed | - | 7 | 20 |

**Table S1: Statistical analysis of the fit results deriving from the comparison of NP multimerization between cell models.** The empirical model $y=1+a\cdot x^k$ ([31]) was used to analyze the data, with the parameter *k* fixed to 0 for HEK and A549 samples (see Figure S1). Resulting values of the fit parameters *a* and *k* are reported along with standard deviations and number of data points (before and after pooling).

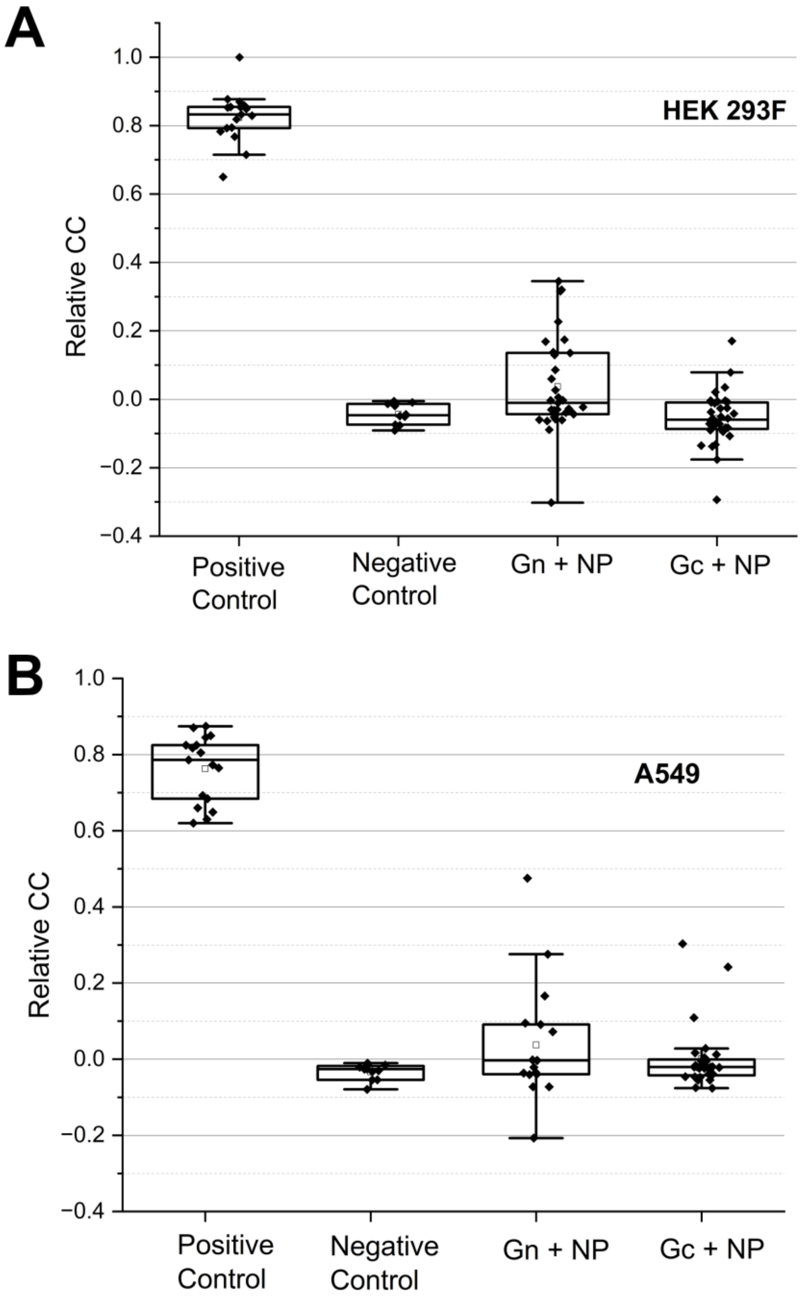


**S2: Average interactions between NP and GPs co-expressed in HEK and A549 cells are weak.** Average relative CC values obtained from [A] HEK cells and [B] A549 cells co-expressing NP and GP (Gn or Gc) using ccN&B. Positive control refers to the tandem cytosolic construct (YFP-mCh2) and the negative control refers to cells co-expressing YFP-NP and cytosolic mCherry2 (mCh2-C1). Each point on the graph represents the average relative CC over a single cell. A minimum of 10 cells are used for each construct combination in every cell model, with a number of independent experiments > 3. The relative CC values calculated here are the pixel average of the ROI selected in each cell (for details, see Materials and Methods)

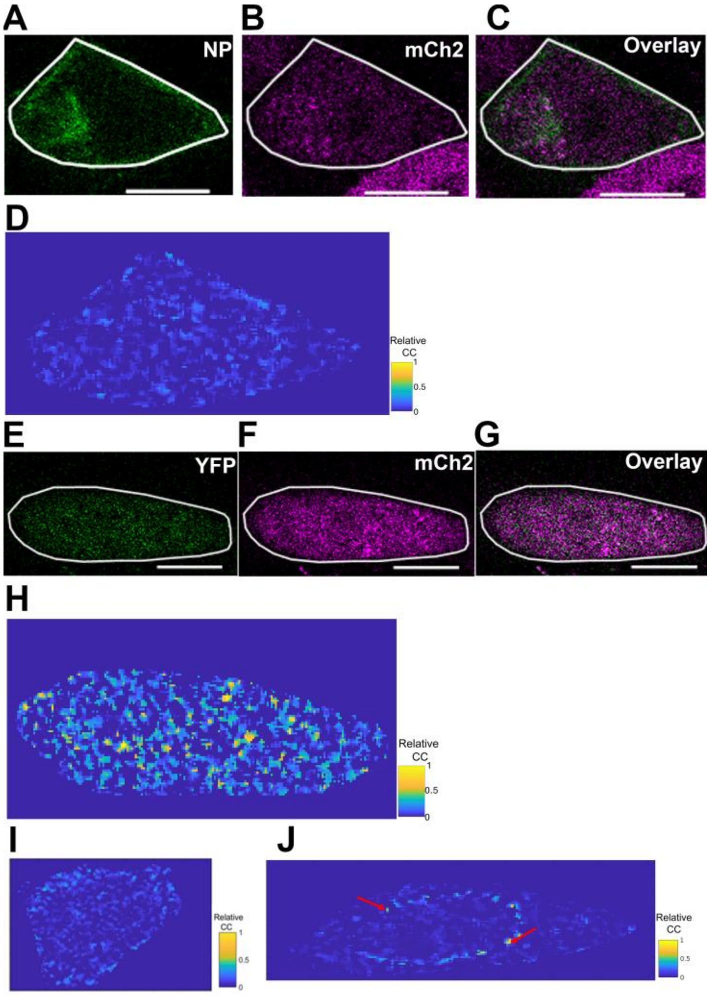


**S3: Intensity and relative CC maps of CHO cells expressing positive and negative control. Panel**s A and B show the YFP and mCh2 channels of confocal images of [A] PUUV YFP-NP and [B] mCh2-C1 (negative control) co-expressed in CHO cells and observed 24 hpt. Panel D shows the relative CC map of the same CHO cell expressing the negative control construct combination. The white line in panels A to C denote the ROI used for the relative CC evaluation, as shown in panel D. Panels E and F show the YFP and mCh2 channels of confocal images of tandem YFP-mCh2 (positive control) expressed in CHO cells and observed 24 hpt. Panel H shows the relative CC map for the same CHO cell. The white line in panels E to G denote the ROI used in the relative CC evaluation, as shown in panel H. Panel I shows an additional representative relative CC image of cells co-expressing NP and Gn. Panel J shows an additional representative relative CC image of cells co-expressing NP and Gc.

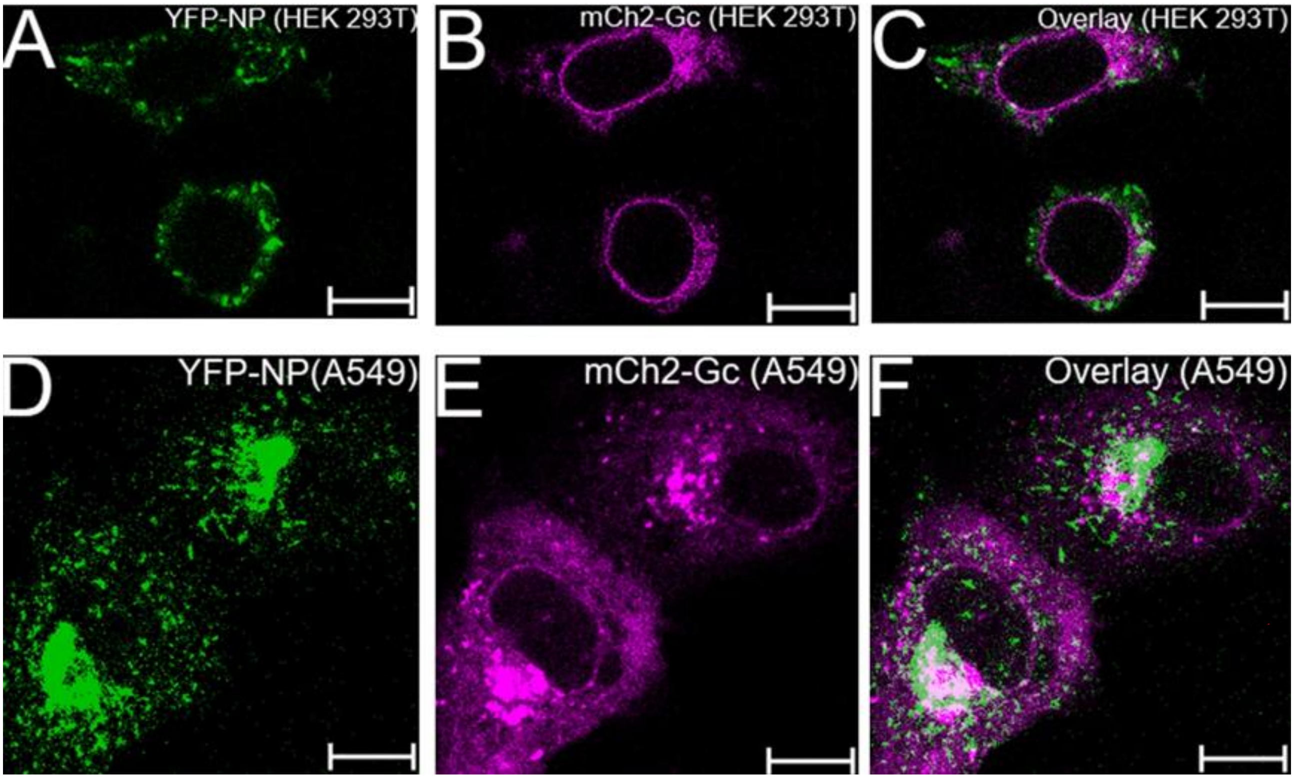


**S4: Co-expression of PUUV NP and PUUV Gc in HEK293T and A549 cells.** Panels A-C show representative confocal images of HEK 293T cells expressing PUUV YFP-NP and PUUV mCh2-Gc. Panel C shows the overlay of the two channels. Panels D-F show representative confocal images of A549 cells expressing PUUV YFP-NP and PUUV mCh2-Gc. Panel F shows the overlay of the two channels. Scale bars are 10 µm.

| Cell Line | a | K | SD (a) | SD (K) | Total N | N pooled |
|---|---|---|---|---|---|---|
| CHO (Gc alone) | 0.54 | 0.7 | 0.23 | 0.6 | 31 | 16 |
| CHO | 1.6 | 0.4 | 0.6 | 0.5 | 25 | 13 |
| HEK | 3.1 | 0.8 | 0.7 | 0.3 | 24 | 12 |
| A549 | 2.0 | 0.8 | 0.7 | 0.7 | 19 | 10 |

| | p-value a | p-value k |
|---|---|---|
| CHO (Gc alone) v/s CHO | <0.01 | 0.44 |
| CHO (Gc alone) vs HEK | <0.01 | 0.99 |
| CHO (Gc alone) vs A549 | <0.01 | 0.84 |
| CHO vs HEK | <0.01 | 0.31 |
| CHO vs A549 | 0.98 | 0.37 |
| HEK vs A549 | 0.02 | 0.82 |

**Table S2: Statistical comparison of fit parameters describing Gc multimerization between cell models co-expressing NP and Gc.** Optimized values of the fit parameters *a* and *k* along with the relevant statistical comparison between different cell models co-expressing NP and Gc are shown in the tables above (see Figure 3 A in the main text). The analytical function used is a non-linear growth equation $1+a(x^k)$, in which *a* and *k* are set as free parameters [31]. The modified Tukey Kramer test was applied for the pairwise comparison. The tables also report the parameters´ standard deviations and number of data points (before and after pooling).

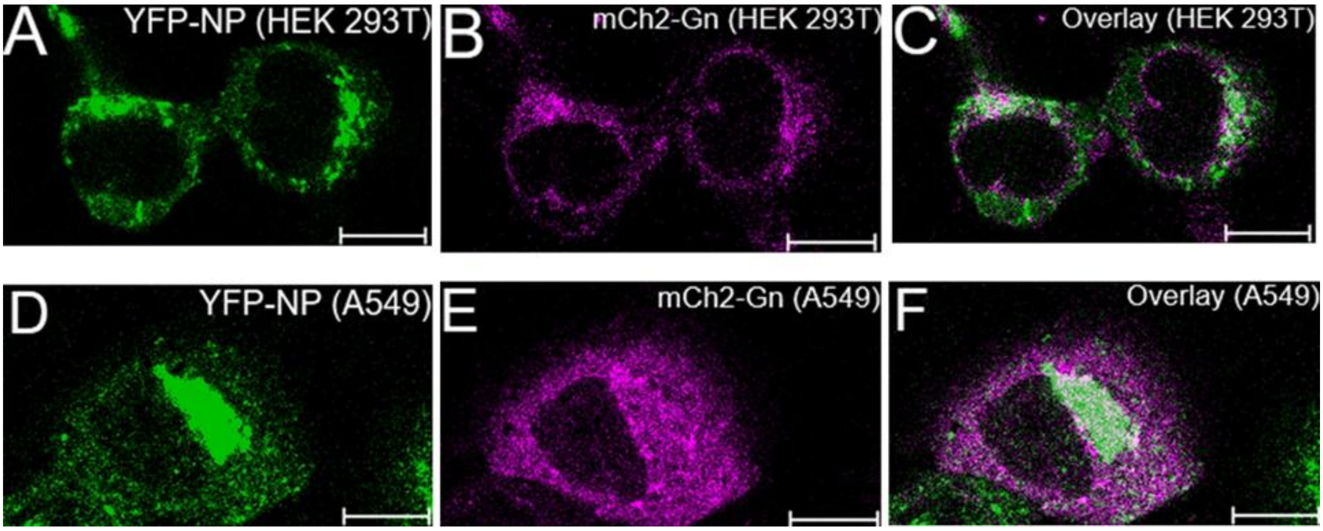


**S5: Co-expression of PUUV NP and PUUV Gn in HEK293T and A549 cells.** Panels A-C show representative confocal images of HEK 293T cells expressing PUUV YFP-NP and PUUV mCh2-Gn. Panel C shows the overlay of the two channels. Panels D-F show representative confocal images of A549 cells expressing PUUV YFP-NP and PUUV mCh2-Gn. Panel F shows the overlay of the two channels. Scale bars are 10 µm.

| Cell Line | $K_4$ | SD | Total N | N pooled |
|---|---|---|---|---|
| CHO (Gn alone) | 3 | 2.8 | 25 | 13 |
| CHO | 4 | 5 | 24 | 12 |
| HEK | 38 | 50 | 20 | 10 |
| A549 | 2.3 | 2.6 | 17 | 9 |

| | p-value, $K_4$ |
|---|---|
| CHO (Gn alone) v/s CHO | 0.8 |
| CHO (Gn alone) vs HEK | 0.2 |
| CHO (Gn alone) vs A549 | 0.7 |
| CHO vs HEK | 0.2 |
| CHO vs A549 | 0.6 |
| HEK vs A549 | 0.2 |

**Table S3: Statistical comparison of fit parameters describing Gn multimerization between cell models co-expressing NP and Gn.** The optimized values of the fit parameter $K_4$, i.e. the association constant for monomer to tetramer equilibrium [32] for PUUV Gn, along with the relevant statistical comparison between different cell models co-expressing NP and Gn are shown in the tables above (see Figure 3 B in the main text). The modified Tukey Kramer test was applied for the statistical pairwise comparison. The tables also report the parameters´ standard deviations and number of data points (before and after pooling).

| Construct | $K_2$ | SD | Total N | N pooled |
|---|---|---|---|---|
| GcΔCT | 0.15 | 0.25 | 18 | 9 |
| GcΔCT (+NP) | 0.5 | 0.7 | 17 | 9 |

| | Parameter | p-value |
|---|---|---|
| GcΔCT vs GcΔCT (NP) | $K_2$ | 0.31 |

**Table S4: Statistical comparison of fit parameter describing GcΔCT multimerization between cell models co-expressing NP and GcΔCT.** The optimized values of the fit parameter $K_2$, i.e. the association constant for monomer to dimer equilibrium [32] for PUUV GcΔCT, along with the relevant statistical comparison between different cell models co-expressing NP and GcΔCT are shown in the tables above (see Figure 4 E in the main text). The modified Tukey Kramer test was applied for the statistical pairwise comparison. The tables also report the parameters´ standard deviations and number of data points (before and after pooling).

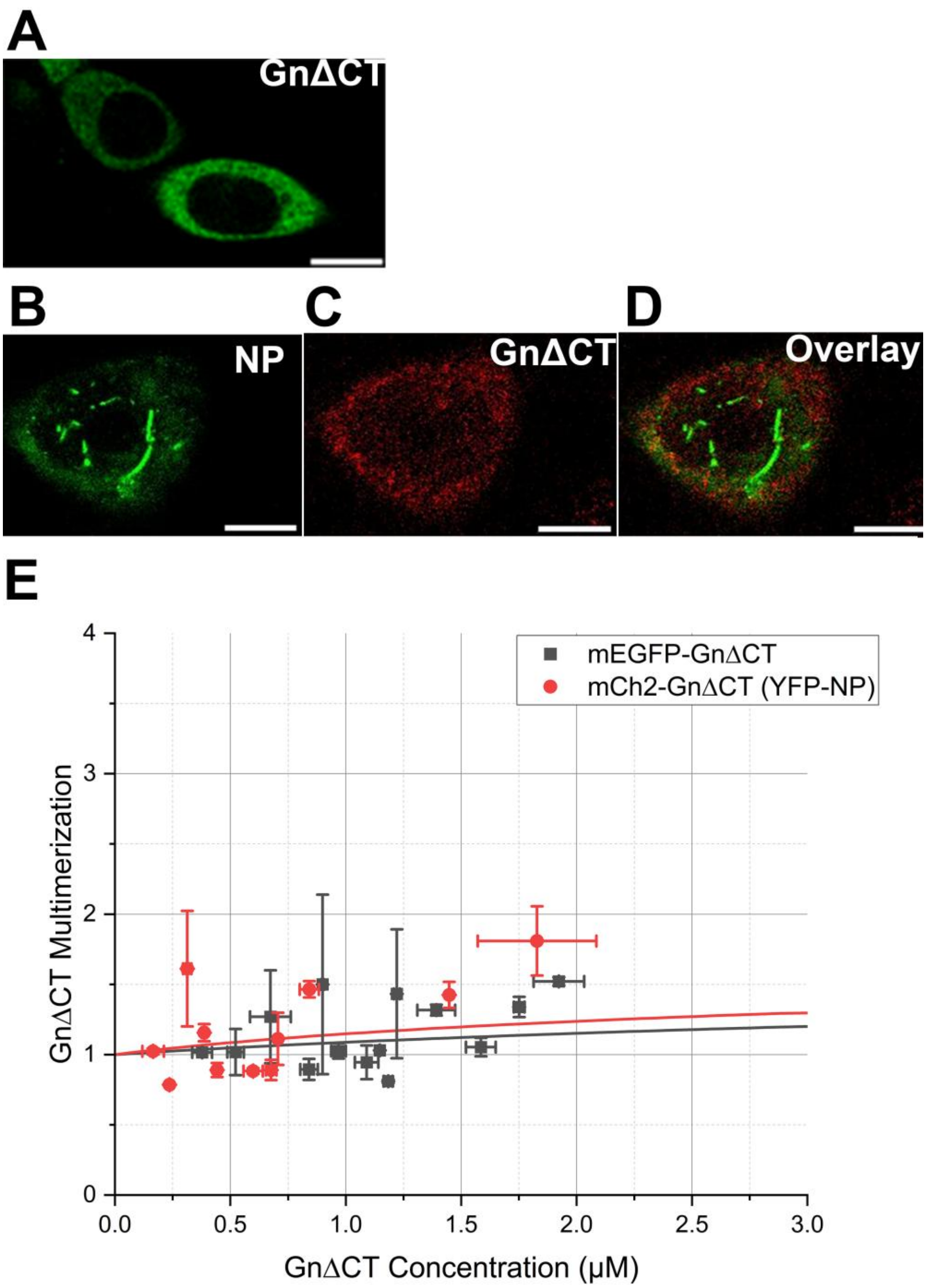


**S6: PUUV GnΔCT remains mostly monomeric, independent of the presence of NP, upon expression in CHO cells.** Panel A shows a representative confocal image of CHO cells expressing PUUV mEGFP- GnΔCT and observed 24 hpt. Panel B - D show typical confocal images of CHO cells co-expressing YFP-NP and mCh2-GnΔCT. Panel E shows the concentration-dependent multimerization analysis of PUUV GcΔCT in the absence (black) and presence (red) of PUUV NP, calculated using N&B analysis. Each point in the graph represents the binned average multimerization values from two cells. The solid lines represent a fit to a monomer-dimer equilibrium model. Scale bars are 10 μm.

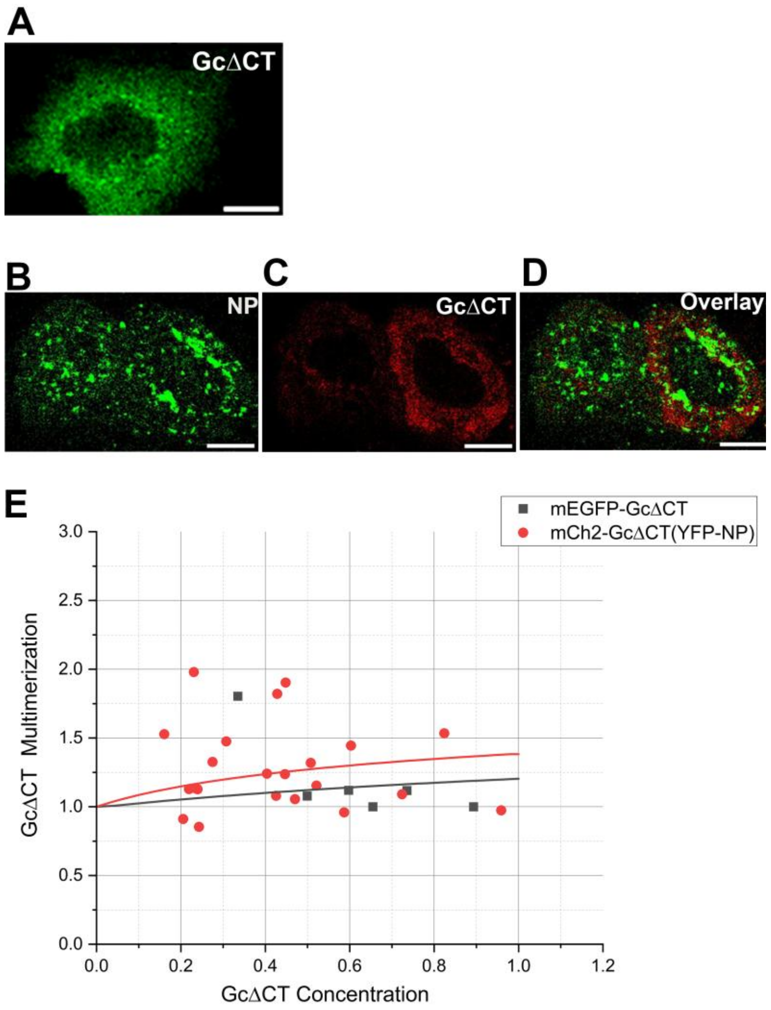


**S7: Lack of GcΔCT inhibits the formation of large Gc assemblies in the presence of NP, also in A549 cells.** Panel A shows a representative image of A549 cells expressing PUUV mEGFP-GcΔCT and observed 24 hpt. Panels B - D show typical confocal images of A549 cells co-expressing YFP-NP and mCh2-GcΔCT. Panel E shows the concentration-dependent multimerization analysis of PUUV GcΔCT in the absence (black) and presence (red) of PUUV NP, calculated using N&B analysis. Each point in the graph represents the binned average multimerization from two cells. The solid lines represent a fit to a monomer-dimer equilibrium model. Fit results and statistical analysis are shown in Table S4. Scale bars are 10 µm.

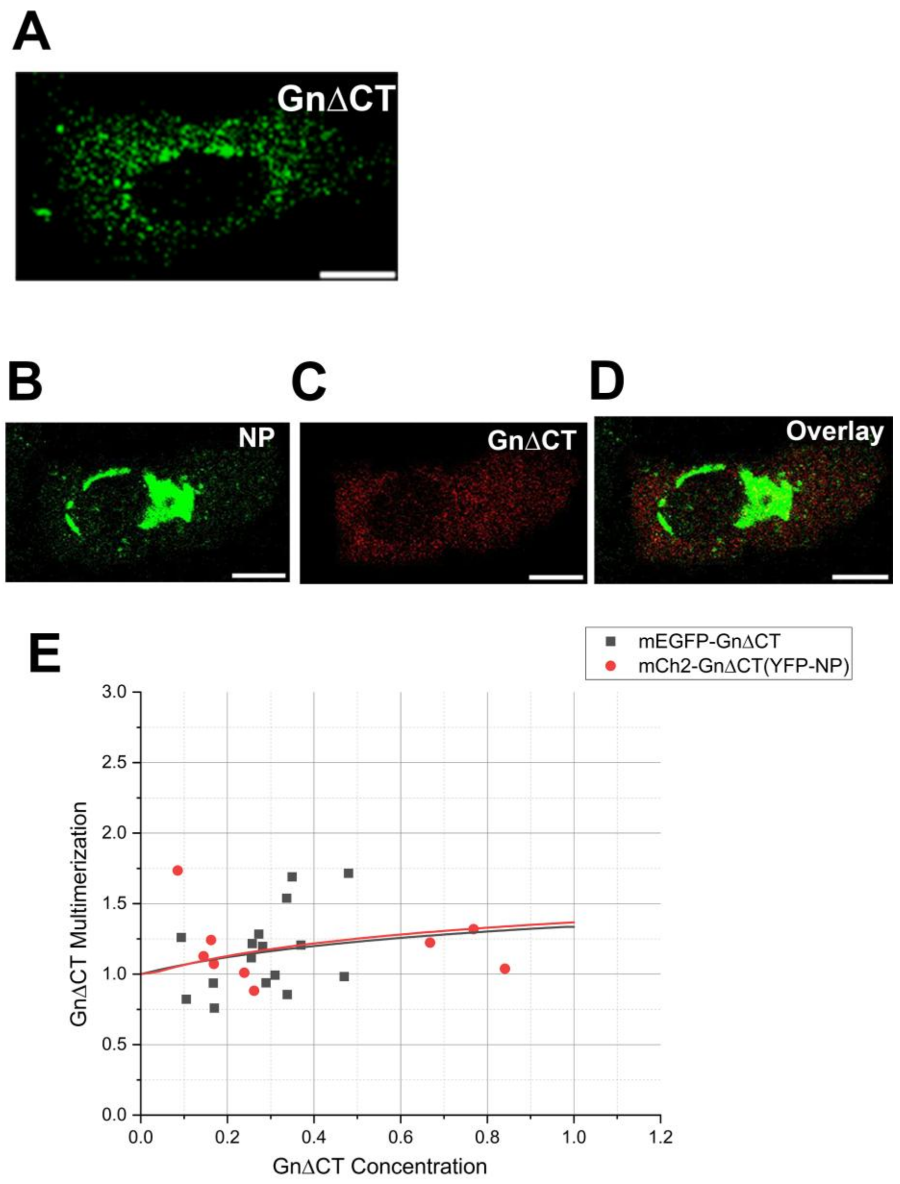


**S8: PUUV GnΔCT is mostly monomeric, independent of the presence of NP, upon expression in A549 cells.** Panel A shows a representative confocal image of A549 cells expressing PUUV mEGFP- GnΔCT and observed 24 hpt. Panel B - D show typical confocal images of A549 cells co-expressing YFP-NP and mCh2-GnΔCT. Panel E shows the concentration-dependent multimerization analysis of PUUV GcΔCT in the absence (black) and presence (red) of PUUV NP, calculated using N&B analysis. Each point in the graph represents the binned average multimerization values from two cells. The solid lines represent a fit to a monomer-dimer equilibrium model. Scale bars are 10 μm.
